# The Sec61/TRAP Translocon Scrambles Lipids

**DOI:** 10.1101/2023.11.23.568215

**Authors:** Matti Javanainen, Sudeep Karki, Dale Tranter, Denys Biriukov, Ville O. Paavilainen

**Affiliations:** Institute of Biotechnology, University of Helsinki, FI-00790 Helsinki, Finland; Central European Institute of Technology, Masaryk University, Kamenice 5, CZ-62500 Brno, Czech Republic

## Abstract

Cell growth relies on the rapid flip–flop of newly synthesized lipids across the ER membrane. This process is facilitated without the need for ATP by specific membrane proteins—scramblases—a few of which have been very recently identified in the ER. We have previously resolved the structure of the translocon-associated protein (TRAP) bound to the Sec61 translocon in the ER membrane, and found this complex to render the membrane locally thinner. Moreover, Sec61 and TRAP each contain a crevice rich in polar residues that can shield a lipid head group as it traverses the hydrophobic membrane environment. We thus hypothesized that both Sec61 and TRAP act as ER scramblases. Here, we characterized the scrambling activity of Sec61 and TRAP using extensive molecular dynamics simulations. We observed that both Sec61 and TRAP efficiently scramble lipids *via* a credit card mechanism. We analyzed the kinetics and thermodynamics of lipid scrambling and demonstrated that local membrane thinning provides a key contribution to scrambling efficiency. Both proteins appear seemingly selective towards phosphatidylcholine lipids over phosphatidylethanolamine and phosphatidylserine, yet this behavior rather reflects the trends observed for these lipids in a protein-free membrane. The identified scrambling pathway in Sec61 structure is physiologically rarely unoccupied due to its role in protein translocation. Furthermore, we found that the scrambling activity of this pathway might be impeded by the presence of ions at a physiological concentration. However, the trimeric bundle of TRAP*β*, TRAP*γ*, and TRAP*δ* might provide scrambling activity insensitive to the functional state of the translocon and the solvent conditions.

## Introduction

Phospholipids are the key building blocks of all cellular membranes. ^1^ Cells thus need to synthesize membrane lipids to support their growth, proliferation, and homeostasis.^2^ These lipids are involved in signaling,^3^ energy storage,^4^ and membrane compartmentalization.^5^ Their majority is synthesized on the cytosolic leaflet of the endoplasmic reticulum (ER) membrane, yet eventually, half of the newly synthesized lipids need to flip to the lumenal leaflet to relieve membrane stress.^6^ In general, lipids show a symmetric distribution across the ER membrane^7^ and are transported to the plasma membrane either along the secretory pathway, through ER–plasma membrane contact sites, or by means of non-vesicular transport.^6^ Phosphatidylserine (PS) is special, as it is exclusively located in the lumenal leaflet of the ER membrane,^7^ and its transport to the plasma membrane relies on dedicated proteins.^7^ These proteins can only pick up lipids from the cytosolic leaflet, indicating that even PS needs to occasionally cross the ER membrane to be transported.^7^

However, phospholipids contain either zwitterionic or anionic head groups, whose spontaneous permeation across the hydrophobic membrane core is energetically unfavorable. Indeed, the measured spontaneous flip–flop half-lifes for zwitterionic phosphatidylcholine (PC) lipids in fluid-phase vesicles is days to weeks. ^8^ Unassisted flip–flops are therefore not likely to form the basis for maintaining lipid symmetry across the leaflets of the ER membrane. This challenge also considers other cellular membranes. The leaflets of the plasma membrane are normally highly asymmetric in composition, ^5,9^ yet in apoptotic cells the distribution of PS becomes symmetric, and its presence in the extracellular leaflet destines the cell for phagocytosis.^10^

Due to the essential biological roles of transbilayer lipid transfer and the associated substantial energetic cost, different membranes contain a suite of dedicated proteins, which significantly enhance the flip–flop rates of phospholipids. Flippases and floppases transfer lipids in specific directions against the concentration gradient by consuming ATP in the process and thus help maintain membrane asymmetry.^11–13^ Scramblases, on the other hand, increase the flip–flop rates bidirectionally without the need for energy input and thus promote leaflet symmetry.^11,12^ The role of proteins in ER membrane scrambling was already acknowledged by the 1980s,^14,15^ yet the identification of these proteins took a further three decades. During that time, it was estimated that lipid transporters constitute 0.2–1.0% of total ER membrane protein, that at least two different transporters are present, and that PC, phosphatidylethanolamine (PE), and PS are all exchanged between leaflets at the same rate, suggestive of using the same transporter proteins.^16,17^

The identification of proteins capable of scrambling lipids without the help of ATP began a decade ago with opsin and TMEM16—two proteins residing in the plasma membrane.^18,19^ Since then, the structure of TMEM16 has also been resolved, ^20^ allowing a description of the scrambling domain formed by hydrophilic residues in the transmembrane domain of TMEM16,^21^ as well as the characterization of its local membrane-thinning ability.^22^ These two features likely contribute to the TMEM16 scramblase activity, and form a basis for our understanding of the structure–function relationship in scrambling.

Very recently, the first ER-resident scramblases—TMEM41B, TME16K, and VMP1— were also identified.^23–28^ Lipid scrambling relies on the ability to transfer polar lipid head groups across the hydrophobic membrane core. From a physical perspective, this challenge resembles that of the integration of membrane proteins with charged residues into the membrane or the translocation of charged proteins through the membrane. Indeed, it has been very recently suggested that lipid scrambling is a general feature of protein insertases and translocases in the ER membrane as well as in mitochondria. ^29^

Recently, we^30^ and other teams^31–33^ resolved the structure of the translocon-associated protein (TRAP) complex associated with the Sec61 translocon in the ER membrane. Sec61 is a gatekeeper for the entry of the vast majority of proteins destined for the secretory pathway. The heterotrimeric Sec61 translocon forms a transmembrane conduit through which secretory proteins enter the ER lumen and membrane proteins get integrated into the ER membrane. Sec61 assembles various ensembles of auxiliary proteins to process different substrates,^33^ with TRAP being required for the translocation of certain proteins such as insulin.^34,35^

Our structural and simulation work revealed that the Sec61/TRAP complex induced local ER membrane thinning.^30^ Moreover, we noticed that the subunits in the TRAP trimeric bundle, namely TRAP*β*, TRAP*δ*, and TRAP*γ*, were also rich in conserved polar amino acids. Together, these subunits also form a polar groove which could potentially facilitate lipid scrambling *via* a similar credit card mechanism as identified for other scramblases.^36,37^ Similarly, in its open conformation, the fairly polar central channel of the Sec61 translocon could be available for the head groups of membrane lipids, although it is presumed to remain closed in the absence of an inserting polypeptide. We thus hypothesized that Sec61 and TRAP act as ER scramblases.

To verify our hypothesis, we characterize here the lipid scrambling by Sec61 and TRAP in detail. We performed extensive coarse-grained simulations of the Sec61/TRAP complex along its various subcomplexes in single- and multi-component lipid membrane environments. We evaluated the ability of these protein complexes to scramble various lipid types that are prevalent in the ER. We identify the scrambling pathways and demonstrate that the membrane-remodeling effect of the Sec61/TRAP complex lowers the free energy barrier for lipid scrambling to a level that is easily crossed at physiological temperature. By performing the simulations at various temperatures, the activation energy for scrambling is obtained through the Arrhenius analysis, and the decomposition of the free energy profiles provides the entropic and enthalpic contributions to the scrambling process. All in all, our results indicate that even if the Sec61 channel is occluded by a polypeptide under translocation, the trimeric TRAP bundle could still provide an efficient pathway for lipid scrambling along with the earlier identified ER scramblases.

## Results and Discussion

### Sec61 and TRAP Both Scramble Lipids

To investigate the ability of Sec61, TRAP, and the Sec61/TRAP complex to scramble lipids, we performed molecular dynamics simulations of these assemblies embedded in a lipid bilayer. For the proteins, we used our recently resolved model for the core Sec61/TRAP complex bound to a mammalian ribosome^30^ based on a combination of cryo-EM and AlphaFold2 prediction.^38^ Since some of the extramembrane parts and flexible loops cannot be unambiguously determined based on cryo-EM in our model (PDB:8BF9), those were built using MODELLER^39^ for this work (see Fig. 1A and B). Notably, our sample contained a substrate-selective KZR-8445 inhibitor, which maintained the Sec61 lateral gate in an open conformation. Therefore, the gate is also open in our model.^40^

**Figure 1:**
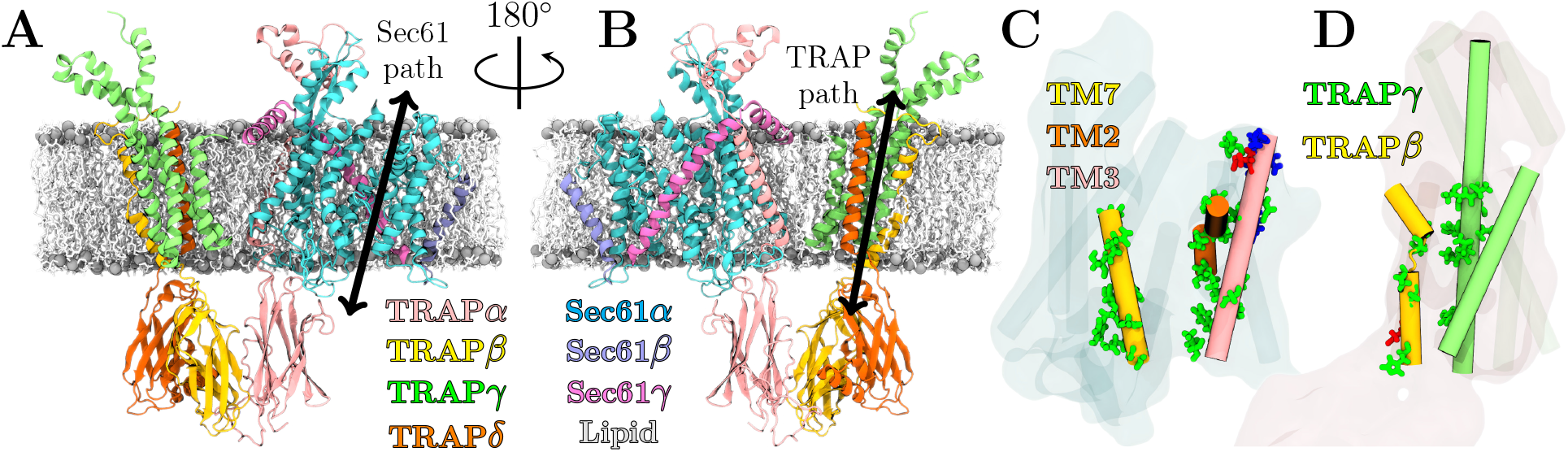
Scrambling pathways of the Sec61/TRAP complex. **A & B)** Snapshots of the Sec61/TRAP complex from A) the side of the lateral gate, “front” and B) from the opposite side, “back”. The prospective scrambling paths of Sec61 along the lateral gate and of TRAP along the groove between the TRAP*β* and TRAP*γ* are highlighted by bidirectional arrows. The structure is embedded in a lipid bilayer shown in gray (phosphorus atoms) and white (rest) to highlight the transmembrane regions. **C & D** The polar (green), anionic (red), and cationic (blue) residues located near the potential scrambling pathways are highlighted for C) Sec61 and D) and TRAP.

The prospective scrambling pathways in Sec61 and TRAP are also highlighted in Fig. 1A and B, respectively. These pathways were predicted based on the presence of groove-like structures containing polar residues that could shield the charged or zwitterionic lipid head groups as they traverse the hydrophobic core of the membrane. The Sec61 pathway is used for the translocation of nascent polypeptides, so the lining of the lateral gate region by polar residues is not surprising. However, the trimeric TRAP bundle also contains a significant amount of polar residues in its transmembrane segments: TRAP*γ* has numerous serines and asparagines in the transmembrane region of its four-helix bundle, whereas TRAP*β* contains a serine and a threonine in the very core of the membrane around its helix-breaking Pro158. These prospective pathways, together with the relevant charged and polar residues, are highlighted in Figs. 1C and 1D for Sec61 and TRAP, respectively.

In order to sample time scales required for spontaneous lipid scrambling, we performed a resolution transformation of our model for the Sec61/TRAP complex into a coarse-grained description following the Martini 3 force field.^41^ We performed simulations with either the complete Sec61/TRAP complex or the isolated trimeric Sec61 or tetrameric TRAP complexes embedded in a 1-palmitoyl-2-oleoyl-*sn*-glycero-3-phosphocholine (POPC) lipid bilayer (Set 1 in Table 1). The functional Sec61/TRAP complex is locked into a specific conformation by ribosomal anchoring,^30^ which we opted to model by restraining the protein backbone in the simulations. This also means that the lateral gate of Sec61—opened by a KZR-8445 inhibitor in our sample^30,40^—remains open throughout the simulations. All simulations, listed in Table 1, were performed in five replicates for 20 µs each using the recommended simulation settings^41,42^ (see Methods for details).

**Table 1:** Simulated systems and the observed flip–flop rates. The rates are reported for different lipid types (*k*_PC_, *k*_PE_, and *k*_PS_), and the overall rate (*k*_tot_) is also provided, all in 1/µs. The single-component membrane was made up of POPC, whereas the multicomponent one consisted of POPC, 1-palmitoyl-2-oleoyl-*sn*-glycero-3-phosphoethanolamine (POPE), and 1-palmitoyl-2-oleoyl-*sn*-glycero-3-phosphoserine (POPS) (see Methods for details). The mean values are calculated from five 20 µs-long replica simulations and shown together with the standard error. With each system here simulated for 5 *×* 20 µs, the total simulation time is *≈*1.5 ms.

| System | $k_{\text{PC}}$ | $k_{\text{PE}}$ | $k_{\text{PS}}$ | $k_{\text{tot}}$ |
| --- | --- | --- | --- | --- |
| **1: Single-component membrane** |  |  |  |  |
| Sec61/TRAP* | $7.7 \pm 0.7$ | – | – | $7.7 \pm 0.7$ |
| Sec61 | $6.5 \pm 1.8$ | – | – | $6.5 \pm 1.8$ |
| TRAP | $2.2 \pm 0.3$ | – | – | $2.2 \pm 0.3$ |
| Sec61/TRAP + NaCl | $1.6 \pm 0.3$ | – | – | $1.6 \pm 0.3$ |
| **2: Multi-component membrane** |  |  |  |  |
| Sec61/TRAP | $2.6 \pm 0.3$ | $0.6 \pm 0.7$ | $1.0 \pm 0.2$ | $4.2 \pm 0.4$ |
| Sec61 | $2.8 \pm 0.4$ | $0.8 \pm 0.2$ | $1.2 \pm 0.5$ | $4.8 \pm 0.8$ |
| TRAP | $0.7 \pm 0.2$ | $0.0 \pm 0.0$ | $0.2 \pm 0.1$ | $1.0 \pm 0.1$ |
| **3: TRAP subunits, multi-component membrane** |  |  |  |  |
| TRAPα | $0.0 \pm 0.0$ | $0.0 \pm 0.0$ | $0.0 \pm 0.0$ | $0.0 \pm 0.0$ |
| TRAPβ | $0.0 \pm 0.0$ | $0.0 \pm 0.0$ | $0.0 \pm 0.0$ | $0.0 \pm 0.0$ |
| TRAPγ | $0.1 \pm 0.1$ | $0.0 \pm 0.0$ | $0.0 \pm 0.0$ | $0.1 \pm 0.1$ |
| TRAPδ | $0.0 \pm 0.0$ | $0.0 \pm 0.0$ | $0.0 \pm 0.0$ | $0.0 \pm 0.0$ |
| **4: Sec61/TRAP complex, varying temperature** |  |  |  |  |
| 290 K | $1.7 \pm 0.4$ | – | – | $1.7 \pm 0.4$ |
| 300 K | $3.3 \pm 0.6$ | – | – | $3.3 \pm 0.6$ |
| 310 K* | $7.7 \pm 0.7$ | – | – | $7.7 \pm 0.7$ |
| 320 K | $14.3 \pm 1.1$ | – | – | $14.3 \pm 1.1$ |
| 330 K | $26.0 \pm 1.0$ | – | – | $26.0 \pm 1.0$ |

\*) Same simulation.

Visual observation of the simulations immediately indicated that lipids were scrambled in the immediate vicinity of the Sec61/TRAP complex, where significant membrane remodeling takes place.^30^ To quantify the scrambling events between the leaflets, we traced the lipid orientations (see Methods). The rates (in lipids/µs) of PC lipids scrambled by the different complexes are listed in the first section of Table 1. From these numbers, it is evident that both Sec61 and TRAP scramble PC lipids at a rate faster than one lipid per microsecond. This corresponds to the *>* 100, 000 lipids scrambled per second *in vitro* by opsin, a known plasma membrane scramblase.^43^ In our simulations, the scrambling of Sec61 is some *≈*3-fold faster than that of TRAP (*p* = 0.0009), yet both contribute at physiologically relevant rates. A similar ratio is also found in the Sec61/TRAP system if the flip–flops are assigned to the two scrambling sites based on distance. The Sec61/TRAP complex scrambles lipids at a rate similar to the sum of the two complexes placed in the membrane alone (*p* = 0.31), indicating that there are no synergistic or antagonistic effects for Sec61 and TRAP. Actually, the scrambling rate of the Sec61/TRAP complex is also not significantly larger than that of Sec61 alone in our simulations (*p* = 0.19).

### Lipids are Scrambled *via* the Credit Card Mechanism

The visual inspection of the simulation trajectories reveals that the flip–flops indeed take place through the crevice formed by either the trimeric TRAP helix bundle or the lateral gate of Sec61. These pathways are also quantified by volumetric densities of the lipid head group beads, extracted from simulations of the Sec61/TRAP complex in a multi-component membrane containing equimolar amounts of POPC, POPE, and POPS (Set 2 in Table 1). These densities are visualized in Figs. 2A and 2B for TRAP and Sec61, respectively. They reveal significant and continuous populations of the lipid head groups in the membrane and between the prospective scrambling pathways shown in Fig. 1C and 1D. The selected events visualized in Fig. 2C and 2D for Sec61 and TRAP demonstrate that lipids are scrambled *via* the credit card mechanism.^36,37^ In this mechanism, the polar head group of the lipid traverses the membrane interior within the polar crevice, whereas its acyl chains remain in the hydrophobic membrane environment. This way, interactions between polar and nonpolar environments are avoided without energy input.

**Figure 2:**
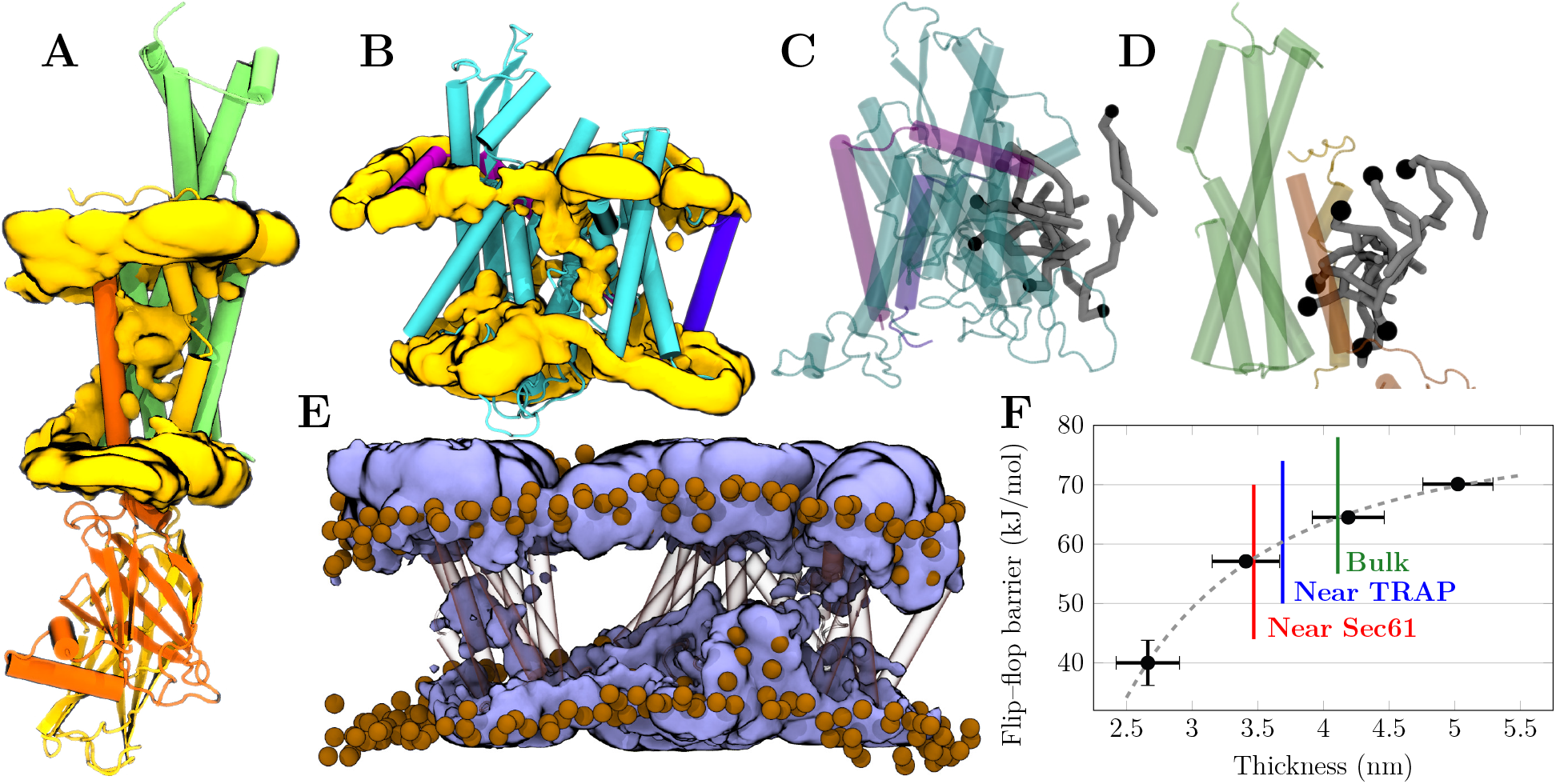
Mechanism of Sec61/TRAP lipid scrambling. **A & B)** Volumetric density maps of the lipid head group phosphate beads (“PO4”) within A) the trimeric bundle of TRAP*β*, TRAP*γ*, and TRAP*δ* subunits or B) Sec61. Averaged from five restrained 20-µs-long simulations of the Sec61/TRAP complex in the multi-component membrane. TRAP and Sec61 subunits are colored as in Fig. 1. The densities of the different lipid types are provided in Figs. S1 (Sec61) and S2 (TRAP) in the SI. For visualization, atomistic protein structures are employed in the rendering. **C & D)** Snapshots of scrambling mechanism by C) the trimeric bundle of TRAP*β*, TRAP*γ*, and TRAP*δ* or D) Sec61. The lipid head group (black bead) partitions to the polar crevices highlighted in Fig. 1C and 1D, whereas the acyl chains (gray) remain in the membrane environment. Coloring of subunits as in Fig. 1 & panels A and B. For visualization, atomistic protein structures are employed in the rendering. **E)** Volumetric density maps of water from atomistic simulations of the Sec61/TRAP complex. Water density is shown in blue, protein in transparent surface with the trimeric bundle of TRAP*β*, TRAP*γ*, and TRAP*δ* on the left and Sec61 with TRAP*α* on the right. Lipid head group phosphorus atoms are shown as brown spheres to highlight membrane thickness and curvature perturbations. **F)** Effect of membrane thickness on the free energy barrier for lipid flip– flop. Black markers show thickness/barrier pairs calculated from a set of bilayers comprised of lipids with saturated chains of varying length. The colored lines show the thickness values observed in the vicinity of Sec61, in the vicinity of the trimeric bundle formed by TRAP*β*, TRAP*γ*, and TRAP*δ*, and far away from the proteins in the “bulk” membrane. The thinning induced by the proteins lowers the free energy barrier by an estimated *≈*4.1 kJ/mol (TRAP) or *≈*6.8 kJ/mol (Sec61). These values are extracted as the differences of the intersections of the colored lines and the gray dashed fit along the *y* axis.

Lipid scrambling by Sec61 is not surprising, considering that the simulations are performed using the open conformation of the Sec61 channel.^30,40^ The open lateral gate provides an extensive trans-bilayer pathway that simultaneously fits multiple lipid head groups and can thus scramble multiple lipids simultaneously, even in opposite directions. This scrambling ability of Sec61 complexed with OST and TRAP was also recently observed in another study using coarse-grained simulations,^29^ and the role of the lateral gate conformation was found to be crucial; with a closed gate, the scrambling ability was largely suppressed. Our analysis indeed found no alternative pathways for scrambling on the Sec61 surface, as evidenced by the lack of continuous inter-leaflet head group densities in Fig. S1B. Still, in our structure, the plug helix and its charged loops somewhat occlude the lateral gate region, and perhaps in another conformation of this region, scrambling could be even faster than observed here.

### TRAP Scrambles More Lipids Under Physiological Conditions

Despite the results presented in the previous sections, it is quite unlikely that Sec61 plays a major role in ER lipid scrambling, as it typically adopts the closed conformation to block undesired calcium leakage. In this conformation, the plug helix and the closed lateral gate seal the lumenal end of the Sec61 channel.^44^ It is only upon the docking of ribosome onto Sec61 that the gate partially opens (“priming”),^45^ whereas the association of the signal peptide renders the gate fully open. ^46^ Thus, Sec61 might not be found in a state with its lateral gate open and not occupied by a nascent chain being translocated by Sec61. Alternatively, the gate can also be opened by Sec61 inhibitors,^40,47^ yet they also associate with the polar residues of the lateral gate along the scrambling pathway. One possibility is that the presence of TRAP and ribosome promotes a more open conformation of the lateral gate and thus assists in initiating the translocation of certain proteins.^30,34^ This could provide a time window for Sec61 to also scramble lipids.

One more reason questions the role of Sec61 as a scramblase under physiological conditions. The simulations described above were performed only with counterions necessary to neutralize excessive protein charges. However, we also repeated the simulation of the Sec61/TRAP complex in the presence of 150 mM of NaCl in the aqueous phase (last entry of Set 1 in Table 1). In five replica simulations, each 20 µs long, the ions crowd the cytosolic vestibule of the Sec61 channel and thereby inhibit lipid scrambling. Indeed, when we assign flip–flops in this system to either the Sec61 or TRAP complex, we find that TRAP scrambles more lipids in the presence of salt (*p* = 0.0005), indicating a drastic change from the *≈*3-fold faster rate observed for Sec61 compared to TRAP observed in the absence of ions. This effect of salt is also corroborated by the volumetric densities of lipid head group beads in Fig. S4, which demonstrate that in the presence of salt, the continuous density bridging the membrane leaflets is lost at the site of Sec61, whereas it is present in the absence of salt. The cytosol is richer in K^+^ than in Na^+^, yet the used Martini 3 model does not provide a model for the former, likely since these two ions are hard to discriminate at the coarsegrained resolution. Still, the observed strong tendency for the monovalent ions to crowd the entry point into the Sec61 channel demonstrates how scrambling can be sensitive to ambient conditions and that Sec61 activity might be especially affected.

These observations contest the role of Sec61 as a physiologically relevant scramblase. Still, TRAP is present in a substantial majority of all ribosome-associated Sec61 complexes,^33^ and it could thus provide a means for lipid scrambling regardless of the functional state of the translocon. Moreover, TRAP scrambling activity is unaffected by the presence of salt, and the scrambling rate of the Sec61/TRAP complex in the presence of salt resembles that of TRAP alone in the absence of salt (Table 1). This is supported by the volumetric density maps in Fig. S4, which demonstrate a continuous density of lipid head groups along the TRAP scrambling pathway regardless of the presence of ions. A plausible explanation for this phenomenon is that, in contrast to Sec61, the transmembrane region of TRAP lacks charged residues, which may attract ions that block the scrambling pathway.

To further look into the structural features behind the observed scrambling ability of TRAP, we also simulated each of the individual TRAP subunits alone (Set 3 in Table 1). In these simulations (5 replicas for each subunit, 20 µs per replica), we only observed a few scrambling events for TRAP*γ*. Moreover, these simulations were performed on a multicomponent membrane containing POPC, POPE, and POPS, indicating that none of these lipid moieties is scrambled efficiently by the individual TRAP subunits. These data indicate that TRAP complex-mediated lipid scrambling arises from cooperation between the subunits. Since the TM domain of TRAP*α* is located far away from the TM domains of other TRAP subunits, it does not contribute to the scrambling ability, and therefore, all of the activity must arise from the six transmembrane domains in the trimeric bundle of TRAP*β*, TRAP*γ*, and TRAP*δ*.

### Polar Intra-Membrane Residues and Membrane Thinning Facilitate Scrambling

Both a membrane-core-spanning pathway of polar residues as well as local membrane thinning have been suggested to contribute to scrambling activity.^21,22^ While we observed lipid scrambling consistently in our simulations, it is still possible that the coarse-grained description omits some key structural or chemical details. Thus, we resorted to all-atom simulations of the Sec61/TRAP complex in a POPC membrane to look into these features. While the structure is independent of the resolution used in the simulations, it is unclear whether the identified scrambling sites (Figs. 1C and 1D) are polar enough to host the lipid head groups. To this end, we calculated the volumetric density map of water molecules from the all-atom simulations. The map in Fig. 2E demonstrates that a significant amount of water penetrates into the membrane interior at both scrambling sites, thus supporting the partitioning of the polar head groups therein. The limited time scale and a more rugged free energy landscape unfortunately preclude the observation of spontaneous lipid flip–flops in atomistic simulations.

Regarding local membrane thinning, our earlier all-atom simulations ^30^ already suggested that the membrane is thinner in the vicinity of Sec61 and TRAP. However, it is unclear how much this thinning contributes to the scrambling activity. To tackle this question, we performed biased accelerated weight histogram (AWH) simulations on a set of PC lipids with saturated acyl chains of increasing lengths (see Methods). We then extracted the free energy barriers and membrane thicknesses from these simulations to estimate the trend between these two quantities as a power law to guide the eye. The data points and this trend are shown in black dots and a gray dashed line, respectively (see Fig. 2E).

Next, we extracted membrane thickness values from the Sec61/TRAP simulations in the multi-component membrane (Set 2 in Table 1). We classified the lipids to be near Sec61 (a head group within 2 nm of any Sec61 subunit) or near TRAP (a head group within 2 nm of the TRAP*β*, TRAP*γ*, or TRAP*δ* subunit), or far from the protein (head group not within 3 nm of any protein subunits). The thicknesses calculated for these lipid populations are shown in Fig. 2E as colored lines. The local thinning due to the presence of the Sec61/TRAP complex seems to contribute to the lowering of the free energy barrier by *≈*7 kJ/mol at most, in case the trend observed for the set of lipids with saturated acyl chains holds. Although it might seem minor in the protein-free case, such a decrease can easily render lipid scrambling thermally accessible if the other mechanism—polar intra-membrane residues—has brought it down to a suitably low level.

### Enthalpic Gain Dominates Over the Loss in Entropy in Scrambling

Next, we estimated the effect of the Sec61, TRAP, and the Sec61/TRAP complex on the scrambling *thermodynamics* by extracting the energetic barriers for lipid flip–flop. For protein-free membranes, we used AWH to extract the potential of mean force (PMF) profile for a POPC flip–flop (see Methods). For the protein-containing system, the scrambling pathway is complicated and thus not easily defined by a single reaction coordinate. Fortunately, the free energy profile can be estimated from lipid head group densities across the membrane due to the spontaneous flip–flops (see Methods). The free energy profiles in Fig. 3A demonstrate that in the absence of proteins, the barrier is *≈*59.2*±*2.2 kJ/mol, and thereby, spontaneous flip–flops are extremely unlikely. In contrast, the presence of Sec61 in the membrane decreases this barrier to *≈*10.8*±*0.5 kJ/mol. With TRAP alone, the barrier decreases to *≈*17.1*±*0.1 kJ/mol, whereas the simultaneous presence of both Sec61 and TRAP results in the smallest barrier of *≈*9.4*±*0.3 kJ/mol for POPC lipids. These numbers follow the scrambling rates in Table 1; the smaller the barrier, the faster the rate.

**Figure 3:**
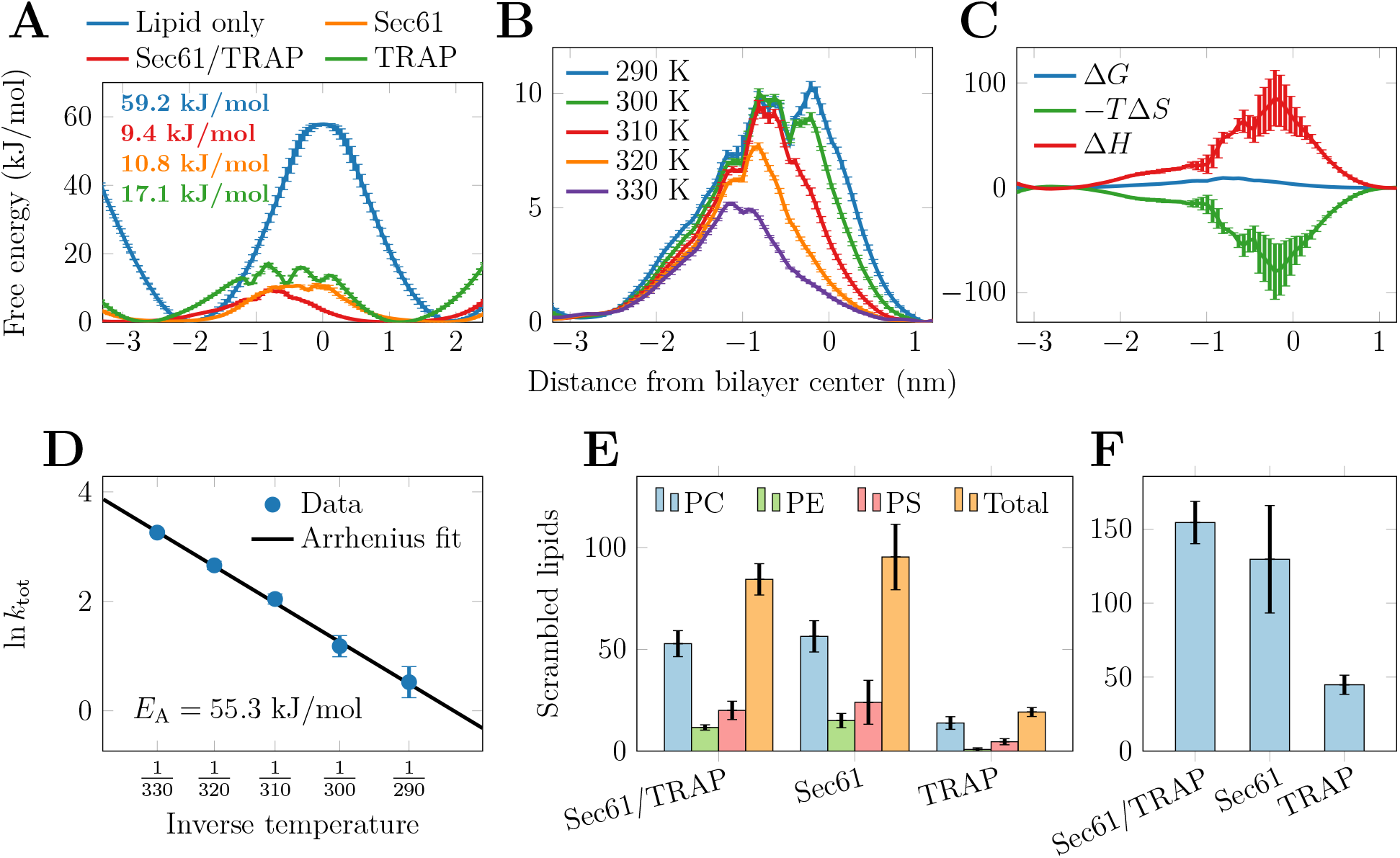
Energetics of lipid scrambling by Sec61/TRAP. **A)** The free energy profiles for a POPC flip–flop in the single-component membrane in the presence of Sec61, TRAP, or the Sec61/TRAP complex as well as in the protein-free membrane at 310 K. The profiles are calculated from density profiles and using biased AWH simulations, respectively (see Methods). Error bars are calculated from the difference of the profiles in the two membrane leaflets (protein-free system) or as the standard deviation of the five replica simulations (protein-containing systems). **B)** Temperature dependence of the free energy profile for lipid scrambling by Sec61/TRAP in the single-component membrane (Set 4 in Table 1). The profiles are calculated from density profiles (see Methods). The error bars show the standard deviation of the five replica simulations. **C)** Decomposition of the free energy profile into entropic and enthalpic components performed by assuming a constant entropy in the simulated temperature range (see Methods). The error bars show the difference between two estimates calculated from two pairs of temperatures (300 K & 320 K as well as 290 K & 330 K, Set 4 in Table 1). **D)** Arrhenius analysis of lipid scrambling rate by Sec61/TRAP in the single-component membranes (Set 4 in Table 1). The scrambling rates observed at different temperatures are fitted with ln *k* = *−E*_A_ *×* 1*/T −* ln *A*, from which the activation energy *E*_A_ is obtained. The error bars show relative error, yet they are not visible for most of the data points. **E)** Lipid head group selectivity of the scrambling activity of Sec61, TRAP, and Sec61/TRAP complexes. Bars show the mean and standard error for the number of total scrambled lipids extracted from the five 20 µs-long replica simulations of the multi-component membrane (Set 2 in Table 1). The membranes contain equal amounts of PC, PE, and PS. **F)** Scrambling of POPC by Sec61, TRAP, and Sec61/TRAP complexes in the single-component membrane (Set 1 in Table 1). Bars show the mean and standard error for the number of total scrambled lipids extracted from the five 20 µs-long replica simulations.

With a simple Eyring model (*k ∼* exp (*−*Δ*G^‡^*)), this difference of *≈*50 kJ/mol results in scrambling a *≈*250 million times faster in the presence of Sec61/TRAP and still *≈*12 million times faster in the presence of TRAP. While in intact phospholipid membranes scrambling takes place in the time scale of a month^8,48^ (half-life), the observed decrease in the energetic barrier would lead to a half-life in the tens of milliseconds for Sec61 and in the hundreds of milliseconds for TRAP. These values are in line with the rates reported for reconstituted VDAC2 dimers.^49^

Simulations also allow us to decompose the free energy profiles into their entropic and enthalpic components. Thus, we performed the simulations with the Sec61/TRAP complex at various temperatures from 290 K to 330 K with 10 K increments (Set 4 in Table 1). The mean free energy profiles for a POPC flip–flop extracted from the five replica simulations performed at each temperature are shown in Fig. 3B. The temperature increase to 330 K leads to the barrier further decrease to *≈*5 kJ/mol. By assuming a constant enthalpy over the used narrow temperature range and thus assigning the observed temperature dependence of the free energy into entropy, the enthalpic and entropic contributions to free energy can be extracted from these simulations (see Methods). The free energy profile, as well as its entropic (*−T* Δ*S*) and enthalpic (Δ*H*) components, are shown in Fig. 3C.

From the profiles in Fig. 3C, it is evident that there is entropic gain in the lipids partitioning to the membrane and being able to sample more conformations than in its canonical membrane orientation. At the free energy barrier, this entropic gain equals *≈*38 kJ/mol, which is approximately half of that calculated for the flip–flop in protein-free membrane (*≈*76 kJ/mol). This is expected, as the credit card mechanism limits the conformations sampled by the lipid significantly in the membrane core as compared to a protein-free flip–flop event. Compared to its canonical positioning in the membrane, however, the acyl chains are looser to sample different configurations in the membrane core, and even the head group can move along the crevice. The drop in entropic gain with the inclusion of the protein indicates that it must come with a substantial gain in enthalpy. Indeed, the presence of the Sec61/TRAP complex can lower the enthalpy value at the free energy barrier for lipid flip–flop from *≈*135 k/mol to *≈*48 kJ/mol due to favorable interactions with intra-membrane polar residues and lipid head groups. It is noteworthy that the maximum (minimum) value of enthalpy (entropy) is larger (smaller) in Fig. 3C than the numbers above, as the latter are reported from the free energy barrier, which does not coincide with the extreme values of the enthalpic and entropic component profiles.

The effect of temperature on scrambling *kinetics* is captured by the Arrhenius formalism, which expects an exponential dependence of the rate on an activation energy *k ∼* exp (*−E*_Arrh_). The natural logarithm of the rates observed in the simulations (in µs*^−^*^1^) as a function of inverse temperature is shown in the Arrhenius plot in Fig. 3D. The measured values nicely fall onto a line, indicating an exponential temperature dependence. Thus, protein-mediated lipid scrambling is well described as an activated process with an activation energy of *≈*55.3 kJ/mol obtained from the fit in Fig. 3D. Unfortunately, few activation energy estimates have been derived from fluid-phase membranes, limiting the comparison of this value with experiments. In DPPC and DMPC vesicles, the values of 122 kJ/mol^8^ and 64 kJ/mol^50^ have been reported, whereas a lower value of 50*±*5 kJ/mol was reported from experiments on supported lipid membranes.^51^ These values highlight that not only the lipid type but also the employed experimental technique could affect the result. Still, the range of experimental values in lipid-only membranes is in the same ballpark as our value for the protein-containing membrane. This indicates that even though the presence of Sec61/TRAP significantly lowers the free energy barrier for lipid flip–flop, it might have a surprisingly little effect on its temperature dependence.

### PC Lipids Are Scrambled Most Efficiently

The currently known lipid scramblases facilitate the bidirectional inter-leaflet movement of lipids regardless of their type—especially head group—yet with varying rates. Earlier experimental work has reported head group-independent scrambling of lipids (PC, PE, and PS) by TMEM14B,^26,27^ whereas TMEM16K was found to scramble PE and PC lipids approximately 3-fold faster than PS.^24^ To study lipid head group preference of Sec61- and TRAP-mediated scrambling, we performed simulations of Sec61, TRAP, and Sec61/TRAP complexes in a larger membrane that contained equimolar amounts of the three major ER lipid moieties: PC, PE, and PS (Set 2 in Table 1). We analyzed the lipid scrambling as earlier, see Table 1 and Fig. 3E.

Unlike for the single-component membranes, where the effect of Sec61 and TRAP were additive, in the multi-component membrane they show a somewhat antagonistic effect; the scrambling rates of the Sec61/TRAP complex are smaller than for the Sec61 and TRAP separately for PC (*p* = 0.005), PE (*p* = 0.039), but not for PS (*p* = 0.14). Regarding lipid-selectivity of the proteins, our analysis revealed that all tested complexes scramble lipids with a high preference for PC. The Sec61 complex scrambles PC more than PE (*p* = 0.0006) and PS (*p <* 0.00001), whereas PS and PE are scrambled at a similar rate (*p* = 0.12). TRAP was also found to be a more efficient scrambler of PC than PE (*p* = 0.00002) or PS (*p* = 0.0003), and the rate for PE was also higher than that for PS (*p* = 0.0012). PS is the only studied lipid with a charged head group, yet its scrambling rates are of the same order of magnitude as those of PC or PE. This suggests that the anionic PS head group does not pose a challenge for the scrambling activity by Sec61 or TRAP, indicating that the polar and well-hydrated crevices along the scrambling path serve as good solvents for the charged phosphate group in the PS head group. These trends are also visible in the volumetric density maps of the PC, PS, and PE head groups that are shown in Fig. S1 and S2 for Sec61 and the trimeric TRAP bundle, respectively. For both scrambling pathways, PC shows the highest occupancy in the membrane core, with PE especially depleted in the case of Sec61.

While it seems at first that the proteins show selectivity towards some lipid types, it is also possible that the observed trends simply follow lipid flip–flop energetics already present in a protein-free membrane. To study this, we again used biased AWH simulations and extracted the barriers for the flip–flops of PC, PE, and PS lipids in a protein-free POPC membrane. The host membrane was always POPC to isolate the effect of the head group of the flipping lipid: a POPE membrane would show tighter packing, thus increasing the free energy barrier, whereas the result in a POPS membrane would likely be affected by counterions. With this approach, we found free energy barriers of 59.2*±*2.2 kJ/mol, 70.6*±*2.4 kJ/mol, and 63.5*±*1.4 kJ/mol, for the flip–flop of PC, PE, and PS, respectively (see Fig. S3 in the SI). This trend follows the observed rates in Fig. 3E, thus suggesting that the proteins are perhaps not selective towards lipid types but instead universally lower the energetic barrier by a similar magnitude, and the differences observed in protein-free membranes are carried over to the protein-containing membranes.

One notable feature in Fig. 3E is that all studied complexes scramble fewer lipids in the multi-component membrane compared to the single-component POPC membrane (Table 1 and Fig. 3F). This likely results from the extended interaction times of the charged PS head group with some polar residues in the scrambling crevices, which can block the scrambling process. This is especially true for TRAP, but the total scrambling rates of Sec61 and the Sec61/TRAP complex are also lower in the multi-component membrane (compare orange bars in Fig. 3E with those in Fig. 3F). Still, the lateral gate crevice in Sec61 is so wide that a single tightly bound PS lipid cannot entirely block the scrambling activity. Similar slower scrambling of PS lipids was also observed experimentally for TMEM16K.^24^

### Our Coarse-Grained Approach Calls for Experimental Validation

The possible methodological limitations of our study are also worth a brief discusssion. The majority of our simulations were performed using a coarse-grained force field, which allows us to obtain reasonable statistics of spontaneous protein-assisted flip–flops. With the time scales reachable with present-day atomistic simulations, this is simply unfeasible to reproduce. Moreover, the free energy landscapes are smoother in coarse-grained models,^52^ lowering the chances of lipids getting stuck in a local minimum along the scrambling pathway. Indeed, in our previous work on the Sec61/TRAP complex, we observed no flip–flops during multiple multi-µs-long all-atom simulations despite observing significant membrane perturbations.^30^ Still, we have obtained information on the hydration of the scrambling path as well as the membrane thinning from atomistic simulations. Another complication is the challenge of balancing the interactions with a limited number of bead types in coarse-grained force fields, which led us to use restrained protein structures. While we used the Martini 3 force field to moderate excessive protein–protein interactions,^41,53^ we still struggled to model the Sec61/TRAP complex without backbone restraints. For Sec61 alone, in simulations with the elastic network and no positional restraints, the Sec61 subunits did not remain correctly associated. For the Sec61/TRAP complex, we attempted to mimic the ribosomal anchoring of Sec61 and TRAP in a specific shape^30^ by restraining only its TRAP*α* and TRAP*γ* subunits proximal to the ribosome in the native structure ^30,32^ (see Methods) and modeling the rest of the protein using elastic network. However, in these simulations, TRAP TM domains collapsed onto the Sec61 structure. This indicates that the protein–protein interactions in the Martini 3 force field might require further tuning, as highlighted by a number of studies;^54–57^ in some cases, these interactions are unable to maintain the subunits together indicating insufficient affinity, whereas in others there is exaggerated interaction. While lipids are still scrambled by these flawed protein conformations, we have not included the related results in the manuscript.

The choice for a criterium used to detect a lipid flip–flop affects the numbers of scrambled lipids, as demonstrated in Fig. S5 in the SI, yet the qualitative trends among the studied complexes were robust to this choice. We attempted to carefully validate the criterium used in this study, yet the observed scrambling rates should still be considered to be ballpark estimates not only due to the challenge of characterizing flip–flops but also due to the inherent limitations of a coarse-grained force field to describe the specific lipid–protein interactions along the scrambling pathway. All in all, the possible limitations of the used coarse-grained models and the methodology involving restraints call for the validation of the observed behavior using experiments, such as *in vitro* scrambling assays.

## Conclusions

Here, we report an extensive set of coarse-grained simulations that collectively demonstrate that both Sec61 and TRAP contain polar crevices that can serve as lipid scrambling pathways in the ER membrane *via* a *credit card mechanism*. While the route along the lateral gate of Sec61 is typically either conformationally inaccessible or likely occluded by an inserting nascent polypeptide, TRAP could serve as a lipid scramblase regardless of the functional state of Sec61.

In addition to containing polar intra-membrane residues, we have earlier demonstrated that Sec61 and TRAP promote local membrane thinning in their vicinity. Our analysis of protein-free membranes suggests that this thinning lowers the free energy barrier of lipid scrambling by up to *≈*7 kJ/mol. Considering that the measured barriers are *≈*10 kJ/mol and *≈*17 kJ/mol for the membranes with Sec61 and TRAP, the effect of thinning plays a significant role in rendering scrambling accessible within thermal fluctuations.

Our free energy decomposition revealed that the presence of the Sec61 and TRAP proteins lowers the free energy barrier by rendering the partitioning of the lipid head groups to the membrane core enthalpically favorable. This is also supported by our atomistic simulations, which highlighted the hydration of the identified scrambling pathways. While the credit card mechanism leads to the loss of conformational entropy compared to a flip–flop in a protein-free system, this is more than compensated by the enthalpic gain.

Finally, we studied the lipid head group selectivity of scrambling by Sec61 and TRAP. In all cases, PC lipids were scrambled most efficiently. Still, our free energy calculations in a protein-free case revealed that the proteins do not show bias towards any head group but rather universally lower the energetic barrier for flip–flop. Still, long residence times of charged PS head groups along the scrambling pathway can lower the overall scrambling rates as compared to PC-only membranes.

Our simulations identify and characterize two novel scrambling pathways in the translocon complex in the ER, where scrambling is essential for membrane growth. Recently, Sec61—along with other insertases—has been suggested to act as a scramblase.^29^ However, the activity of Sec61 heavily depends on the conformation of its lateral gate and the occupancy of the open conformation with an empty Sec61 channel is expected to be low. Moreover, our simulations revealed that the scrambling by Sec61 is sensitive to the ionic environment, as the inclusion of NaCl salt led to a significant reduction in the scrambling rate. A detailed analysis revealed that in this case, scrambling takes almost exclusively place along the pathway on the TRAP trimeric bundle. Therefore TRAP—as an almost stoichiometric partner of Sec61 in the translocon—could provide a more efficient scrambling pathway available in a physiological setting despite the functional state of the translocon. These findings naturally call for experimental validation, yet Sec61 is notoriously challenging to purify and reconstitute,^58^ let alone complexed with TRAP.

## Experimental

### Coarse-Grained Simulations

#### Simulation Systems

We generated two sizes of membranes containing different protein complexes: Sec61, TRAP, Sec61/TRAP, TRAP*α*, TRAP*β*, TRAP*γ*, or TRAP*δ*. First, Sec61, TRAP, or the Sec61/TRAP were embedded in smaller single-component membranes made up of POPC (Set 1 in Table 1). These membranes contained 600 POPC lipids, 40 water beads per lipid (total of 25000), and counter ions (Cl*^−^* or Na^+^) to neutralize the excess protein charge. Additionally, the mem-brane with Sec61/TRAP was also simulated in the presence of *≈*150 mM of NaCl.

Secondly, Sec61, TRAP, or the Sec61/TRAP were also embedded in larger multi-component membranes whose leaflets both originally contained 900 lipids with phosphatidylcholine (PC), phosphatidylethanolamine (PE), and phosphatidylserine (PS) lipids present at equal amounts (Set 2 in Table 1). The choice not to mimic the exact composition of the ER mem-brane^1^ was justified on the proper sampling of the flip–flop events of these different lipid types in the simulation time scale; very few lipids might not diffuse to the protein complex often enough for flip–flops to happen at statistically significant amounts. The membranes were solvated by 35 water beads per lipid for a total of 63000 beads. 600 Na^+^ ions were included to neutralize the charges of the PS head groups, and additional Na^+^ or Cl*^−^* ions were included to neutralize the excess charge of the protein complex present. These larger multi-component membranes were also simulated in the presence of individual TRAP subunits (Set 3 in Table 1). In all simulated membranes, the lipids were modeled to contain palmitate and oleate acyl chains.

The systems were modeled using the latest version 3 of the Martini force field.^41^ The systems were generated using CHARMM-GUI and equilibrated following the CHARMM-GUI protocol.^59^ Then, a 1 µs-long simulation was performed in which the membrane temperature was set to 400 K for fast lipid mixing. The lipid head groups were restrained in the direction normal to the membrane to prevent any flip–flops at this stage. Five frames, separated by 200 ns of simulation time, were extracted and used as initial configurations for the five 20 µs-long production simulations at 310 K. In these simulations, the backbone of the entire protein complex was restrained.

The Sec61/TRAP complex in the smaller single-component POPC membrane was also simulated at multiple temperatures, namely 290, 300, 310, 320, and 330 K (Set 4 in Table 1). Again, five 20 µs-long replicas were performed at each temperature. Here, the goal was to study the activation energy as well as the free energy components of the flipping process, and a smaller membrane with less fluctuations resulted in a more well-defined density profile from which the free energy profile was estimated.

Additionally, we performed simulations of the Sec61/TRAP system with the TRAP*α* and TRAP*γ* subunits restrained only at the ribosomal anchoring sites, namely between W255 and K266 of TRAP*α* as well as between R110 and K115 of TRAP*γ*, constituting the TRAP*α* anchor and TRAP*γ*finger, respectively.^32^ The Sec61 simulation was also repeated without any restraints. The elastic network approach was used in simulations in which the entire protein was not restrained to maintain the tertiary structure. As explained in the Results section, data extracted from these simulations were eventually not used in any figures.

The free energy barriers for lipid flip–flop in protein-free membranes were calculated using a simulation system containing a total of 201 lipids (100 per leaflet + one that was biased to perform a flip–flop). We performed such simulations for 1) a series of PC phospholipids with saturated acyl chains of different lengths, DPTC (2 beads per acyl chain), DLPC (3 beads), DPPC (4 beads), and DBPC (5 beads); 2) a series of lipids with palmitoyl and oleoyl acyl chains and PC, PE, or PS head groups. In the latter case, the membrane consisted of POPC lipids, and only the identity of the flipping lipid varied (POPC, POPE, or POPS).

#### Simulation Parameters

The simulations were performed using the SYCL implementation of GROMACS v. 2023 on AMD GPUs.^60,61^ The equations of motion were integrated with the leap-frog integrator with a time step of 25 fs. Buffered Verlet lists were used to keep track of neighboring beads.^62^ Reaction field electrostatics, with a dielectric constant of 15 within the cutoff of 1.1 nm and *∞* beyond it, was used, following the recommended parameters.^42^ The Lennard-Jones potential was shifted to zero at a distance of 1.1 nm. The temperatures of the protein, lipids, and solvent were separately maintained at 310 K with the stochastic velocity rescaling thermostat^63^ with a time constant of 1 ps. The pressure was maintained at 1 bar using the Parrinello– Rahman barostat^64^ with semi-isotropic coupling (two dimensions along the membrane plane coupled together), a time constant of 12 ps, and a compressibility of 3*×*10*^−^*^4^ 1/bar.

The free energy barrier for lipid flip–flop in the protein-free membranes was extracted using the accelerated weight histogram (AWH) technique.^65^ Apart from AWH settings, we used identical simulation parameters as for the protein-containing simulations. For AWH, we set the reaction coordinate to be the distance between the head group phosphate bead of a single lipid and the rest of the lipid bilayer. The distance was measured along the *z* axis, *i.e.*, along the direction normal to the bilayer. The range of [-4.5,4.5] of this reaction coordinate was sampled, which resulted in the lipid performing multiple flip–flops during the AWH simulation. When it comes to the AWH options, we used the geometry direction. The convolved potential shape was used, and the two-stage approach was used for rapid convergence. The force constant was set to 10^5^ kJ/(mol*×*nm^2^). The target coordinate distribution was set to uniform (“constant”). All AWH simulations were 5 µs long.

#### Simulation Analyses

##### Flip–flop Detection

The flip–flops were characterized by events when a lipid changed its orientation from that characteristic of the upper leaflet to that characteristic of the lower leaflet or *vice versa*. These orientations were defined by a vector connecting the phosphate bead (“PO4”) and the last bead of one of the acyl chains (“C4B”); if the *z* coordinate of the PO4 bead was 2.1 nm smaller (larger) than that of C4B, the lipid was assigned to the upper (lower) leaflet. The numbers of flip–flops visually counted in simulations with individual TRAP subunits (such a visual observation was possible for these systems due to their small number of flip–flops) were reproduced with threshold values of 2.1–2.4 nm. The flip–flop numbers in the single-component membranes with Sec61 or TRAP (Set 1 in Table 1) were also found to plateau around these values, as demonstrated in the top panel of Fig. S5. Moreover, the difference between Sec61 and TRAP scramblase activity was found to be maximal at the lower end of the range (bottom panel of Fig. S5). Thus, we used a threshold value of 2.1 nm in all analyses. Events where a lipid changed its assigned leaflet from one another were recorded as flip–flops.

In the Sec61/TRAP systems, flip–flops were assigned to the protein complex (Sec61 or TRAP) if it initiated within 2 nm of the corresponding scrambling pathway.

##### Activation Energies

The activation energies (*E_A_*) for the flip–flop process were extracted with Arrhenius analysis from the flip–flop rates (*k*_flip_*_−_*_flop_) as

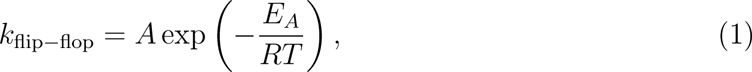

where *A* is a temperature-independent prefactor, *R* is the universal gas constant, and *T* is temperature.

##### Free Energy Profiles from Unbiased Simulations

The free energy profiles for lipid flip–flop in protein-containing systems were estimated from the density profiles of the phosphate (“PO4”) bead as Δ*G* = *−RT* ln(*ρ*(*z*)*/ρ*_0_), where *ρ*(*z*) is the local density along the *z* axis (normal to the membrane), and *ρ*_0_ an arbitrary scaling factor chosen so that Δ*G* = 0 in the equilibrium position of PO4 in the membrane. For the density profiles, the system was divided into 300 slices. Since the membrane shape fluctuates, we set a reference to the protein frame. For the trimeric TRAP bundle, we used residues Thr149 to Ala173 of TRAP*δ*, *i.e.* its transmembrane helix. For Sec61, we used the helical region of TM3, spanning from Lys97 to Met123. For the Sec61/TRAP complex, we used the union of these two groups.

##### Free Energy Decomposition

The entropic and enthalpic components of the flip–flop process were evaluated assuming a constant entropy component between two pairs of temperatures: 300 K and 320 K (Δ*T* = 20 *K*) as well as 290 K and 330 K (Δ*T* = 40 *K*) using^66^

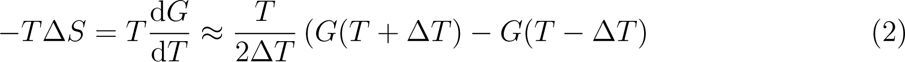

and

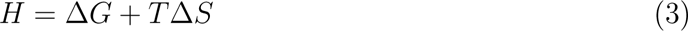

##### Membrane Thickness

Membrane thickness values were calculated using g lomepro^67^ and based on the local inter-leaflet distance of the phosphate (“PO4”) beads.

### All-Atom Simulations

We also simulated the Sec61/TRAP complex embedded in a POPC membrane using the all-atom CHARMM36 force field.^68^ The membrane consisted of 400 POPC lipids and was solvated with 100 water molecules per lipid (total of 40000). The excess protein charge was neutralized by Na^+^ ions.

The atomistic system was simulated for 300 ns with an integration time step of 2 fs. Buffered Verlet lists were used to keep track of atomic neighbors.^62^ Long-range electrostatics were implemented using the smooth PME approach. ^69,70^ The Lennard-Jones potential was cut off at 1.2 nm, and the forces were switched to zero between 1.0 and 1.2 nm. The temperatures of the membrane components (protein and lipids) and the solvent were separately maintained at 310 K using the Nośe–Hoover thermostat^71,72^ with a time constant of 1 ps. The pressures normal to the membrane plane (*z*) and along it (*x* & *y*) were maintained at 1 bar using the Parrinello–Rahman^64^ barostat with semi-isotropic coupling. The time constant of the barostat was set to 5 ps and the compressibility to 4.5*×*10*^−^*^5^ bar*^−^*^1^. Bonds involving hydrogens were constrained using P-LINCS,^73,74^ and water geometry was constrained using SETTLE.^75^

## Data availability

All coarse-grained simulation data with protein-containing systems are available online in the Zenodo repository at DOIs: 10.5281/zenodo.10166590, 10.5281/zenodo.10168857, and 10.5281/zenodo.10169434.

## Acknowledgement

We thank the Research Council of Finland (grant nos. 338836 and 314672 for VOP and grant no. 338160 for MJ) and the National Institutes of Health (grant NIGMS R01GM132649 for VOP) for funding as well as CSC–IT Center for Science (Espoo, Finland) for computational resources. DB acknowledges support from the project “National Institute of Virology and Bacteriology (Program EXCELES, ID Project No. LX22NPO5103) – Funded by the European Union – Next Generation EU”.

## Supporting Information Available

Volumetric maps of lipid densities along the scrambling paths, including for the simulation with salt. Free energy profiles for PC, PE, and PS flip–flop in a POPC membrane.

## Supplementary Information

**Figure S1:**
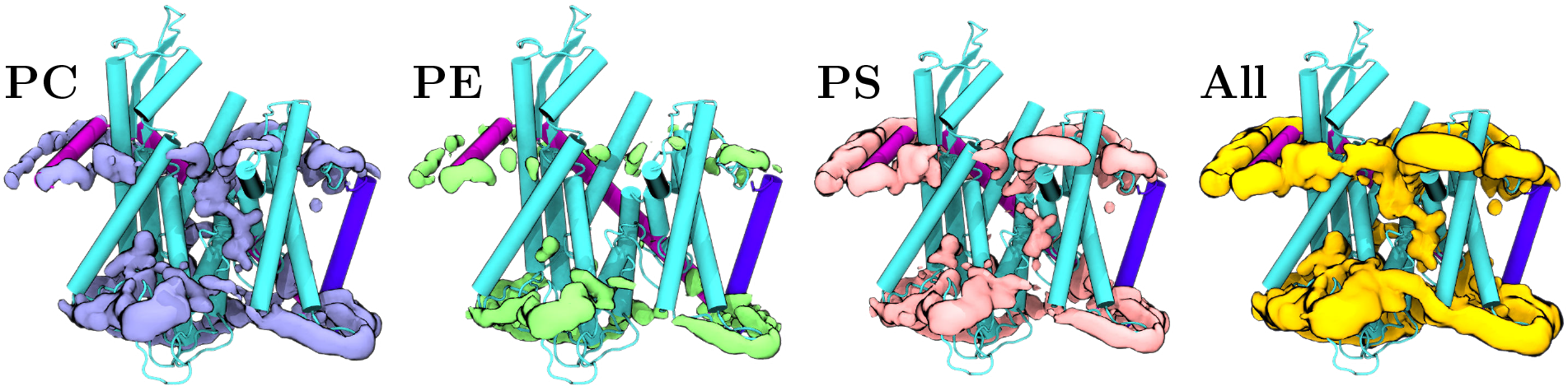
Volumetric maps of the densities of lipid moieties around Sec61. The maps are calculated from the simulation with the Sec61/TRAP complex present in the three-component lipid membrane. The densities are shown separately for all components, and the map for the total density (“All”) is also repeated from the main text for easy comparison. The all-atom structure for the protein is used for visualization purposes. Coloring as in Fig. 1: Sec61*α* is shown in cyan, Sec61*β* in blue, and Sec61*γ* in purple. Lipids are colored as in Fig. 3E. The threshold for the maps was set at 0.01 Å*^−^*^3^ in VMD, and all replicas were used in the analysis (5 *×* 20 µs).

**Figure S2:**
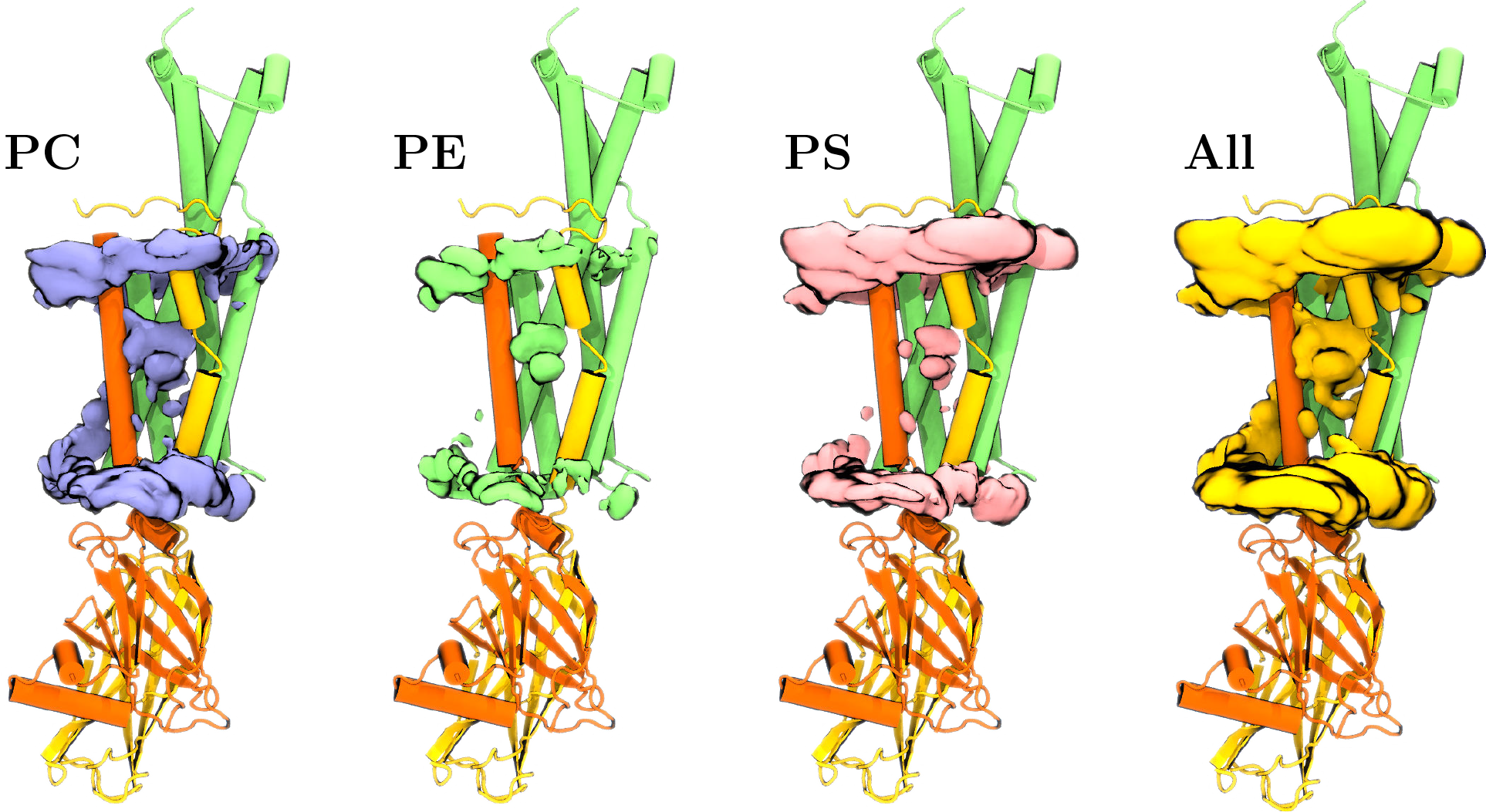
Volumetric maps of the densities of lipid moieties around the bundle of TRAP*β*, TRAP*γ*, and TRAP*δ*. The maps are calculated from the simulation with the Sec61/TRAP complex present in the three-component lipid membrane. The densities are shown separately for all components, and the map for the total density (“All”) is also repeated from the main text for easy comparison. The all-atom structure for the protein is used for visualization purposes. Coloring as in Fig. 1: TRAP*β* is shown in yellow, TRAP*γ* in green, and TRAP*δ* in orange. Lipids are colored as in Fig. 3E. The threshold for the maps was set at 0.01 Å*^−^*^3^ in VMD, and all replicas were used in the analysis (5 *×* 20 µs).

**Figure S3:**
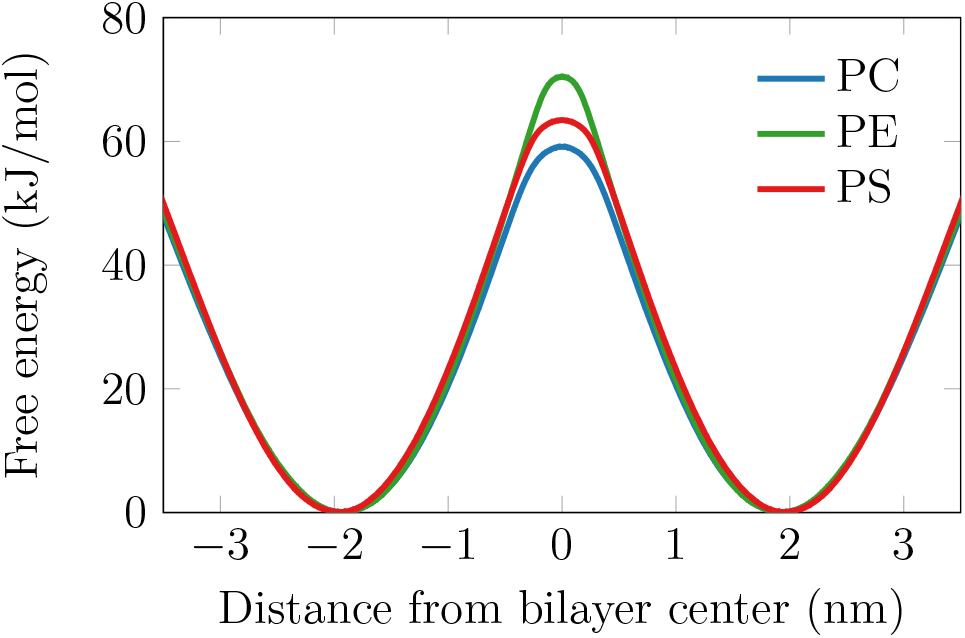
Free energy profiles for lipid flip–flop in a POPC membrane. The profiles are extracted for palmitoyloleoylphosphatidylcholine lipids with three different head groups: PE, PC, and PS. The host membrane is POPC. The profiles obtained from the AWH method are symmetrized, and the AWH error estimates are 0.26–0.37 kJ/mol.

**Figure S4:**
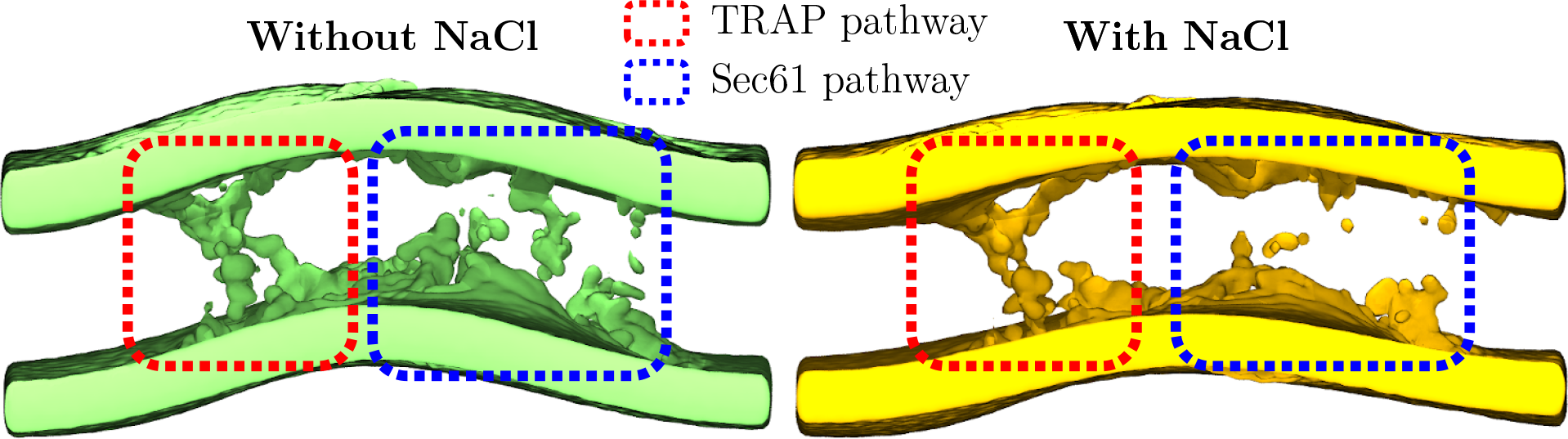
Effect of NaCl salt on lipid scrambling. Data are shown for the single-component POPC membrane. The surface presentation shows the volumetric density of the phosphate (PO4) bead of POPC. The threshold for the maps was set at 0.02 Å*^−^*^3^ in VMD. The two panels show the data for the system without NaCl (counter-ions to neutralize the charge of protein residues are still included) and with *≈*150 mM NaCl. The two pathways are highlighted with colored rectangles. In the presence of salt, the Sec61 pathway is not accessible to the PC head group, whereas the TRAP pathway is unaffected.

**Figure S5:**
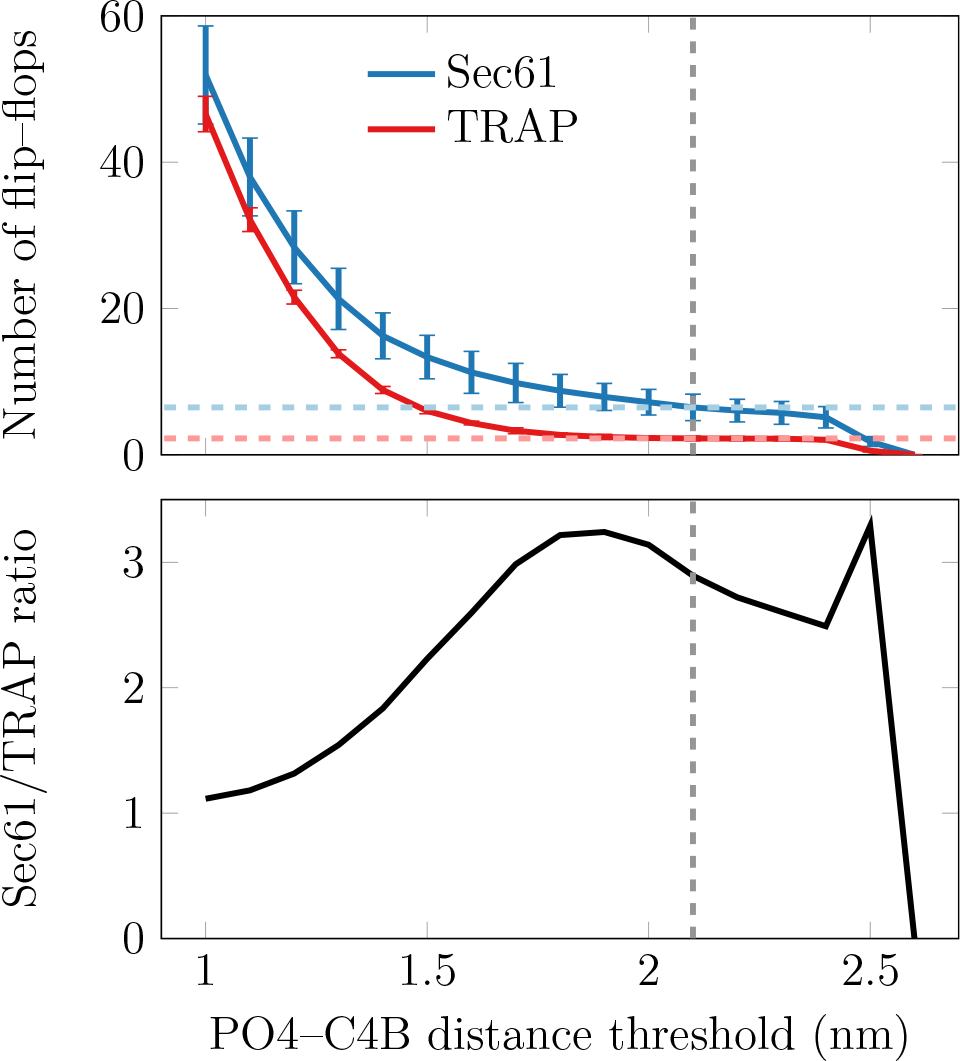
Effect of detection threshold on the number of observed flip–flops. Top: The numbers of flip–flops in the single-component membranes (Set 1 in Table 1) for Sec61 and TRAP as a function of PO4–C4B distance threshold used in flip–flop detection. The dashed gray lines show the value of 2.1 nm used in all calculations. The light blue and light red lines highlight that the number of flip–flops is relatively insensitive around the chosen value. Bottom: The ratio of flip–flops in the systems with Sec61 and TRAP. The gray dashed line again shows the chosen value of 2.1 nm.

